# CRISPR-Cas9 genome editing in human cells works via the Fanconi Anemia pathway

**DOI:** 10.1101/136028

**Authors:** Chris D Richardson, Katelynn R Kazane, Sharon J Feng, Nicholas L Bray, Axel J Schäfer, Stephen Floor, Jacob E Corn

## Abstract

CRISPR-Cas9 genome editing creates targeted double strand breaks (DSBs) in eukaryotic cells that are processed by cellular DNA repair pathways. Co-administration of single stranded oligonucleotide donor DNA (ssODN) during editing can result in high-efficiency (>20%) incorporation of ssODN sequences into the break site. This process is commonly referred to as homology directed repair (HDR) and here referred to as single stranded template repair (SSTR) to distinguish it from repair using a double stranded DNA donor (dsDonor). The high efficacy of SSTR makes it a promising avenue for the treatment of genetic diseases^1,2^, but the genetic basis of SSTR editing is still unclear, leaving its use a mostly empiric process. To determine the pathways underlying SSTR in human cells, we developed a coupled knockdown-editing screening system capable of interrogating multiple editing outcomes in the context of thousands of individual gene knockdowns. Unexpectedly, we found that SSTR requires multiple components of the Fanconi Anemia (FA) repair pathway, but does not require Rad51-mediated homologous recombination, distinguishing SSTR from repair using dsDonors. Knockdown of FA genes impacts SSTR without altering break repair by non-homologous end joining (NHEJ) in multiple human cell lines and in neonatal dermal fibroblasts. Our results establish an unanticipated and central role for the FA pathway in templated repair from single stranded DNA by human cells. Therapeutic genome editing has been proposed to treat genetic disorders caused by deficiencies in DNA repair, including Fanconi Anemia. Our data imply that patient genotype and/or transcriptome profoundly impact the effectiveness of gene editing treatments and that adjuvant treatments to bias cells towards FA repair pathways could have considerable therapeutic value.

## Main Text

The type II CRISPR endonuclease Cas9 and engineered guide RNA (gRNA) form a ribonucleoprotein (RNP) complex that introduces double stranded breaks (DSBs) at DNA sequences complementary to the 23 bp protospacer-PAM sequence. This activity stimulates two major types of DNA repair within a host cell that are relevant to genome editing: genetic disruption, which creates insertions or deletions (indels) at the cut site and can disrupt functional sequences; and genetic replacement, which incorporates exogenous donor DNA sequences at the cut site, allowing the correction of dysfunctional elements or insertion of new information^3^. Efficient and targeted genetic replacement is particularly exciting, as it holds great promise for the cure of myriad genetic diseases.

Despite the rapid adoption of CRISPR-Cas9 genome editing, relatively little is known about which cellular DSB repair pathways underlie Cas9-mediated genetic replacement. This lack of clarity has complicated efforts to better understand and rationally improve the process of genome editing. The pathways responsible for genetic replacement are frequently referred to in aggregate as HDR, which includes DSB repair programmed from dsDonors (both linear and plasmid) requiring several kilobases of homology to the targeted site, as well as synthetic ssODNs with only 100-200 bases of homology to the target^4^. Repair from dsDonors is relatively inefficient in most cell types^5^ and is assumed to utilize a repair mechanism paralleling meiotic homologous recombination (HR)^6^. By contrast, SSTR is highly effective in human cells (>20% of alleles)^1,5,7^ and broadly conserved among metazoans^8^, but very little is known about the mechanism responsible. While screening human cancer cell lines, we found that SSTR-based genome editing at a given locus can vary from completely ineffective (0% SSTR) to extremely efficient (30% SSTR) depending on the cell background [**Extended Data Figure 1**]. This implies genetic or transcriptional differences that up- or down-regulate gene editing in different contexts.

To map the pathways involved in SSTR, we developed a coupled inhibition-editing screening platform that combines individual CRISPR inhibition (CRISPRi) of thousands of genes with Cas9 editing at a single-copy genomically integrated BFP reporter **[Figure 1A].** Each cell in the screening pool stably expresses a dCas9-KRAB CRISPRi construct as well as a gRNA targeting the TSS of a single gene. This pool is then nucleofected with preformed Cas9-gRNA ribonucleoprotein complex (RNP) targeting the BFP reporter, as well as an ssODN that programs a 3 basepair codon-swap that converts BFP to GFP^7^. The femtomolar affinity between *S. pyogenes* Cas9 and the gRNA^9^, along with the transient nature of the Cas9 RNP^10^ strongly disfavors guide swapping between Cas9 molecules, preserving separation between CRISPRi and targeted gene editing. Editing outcomes in each cell are separated by fluorescence activated cell sorting (FACS) and next generation sequencing is used to determine genes whose knockdown leads to enrichment or depletion from each sorted population. [**Figure 1A]**.

**Figure 1:**
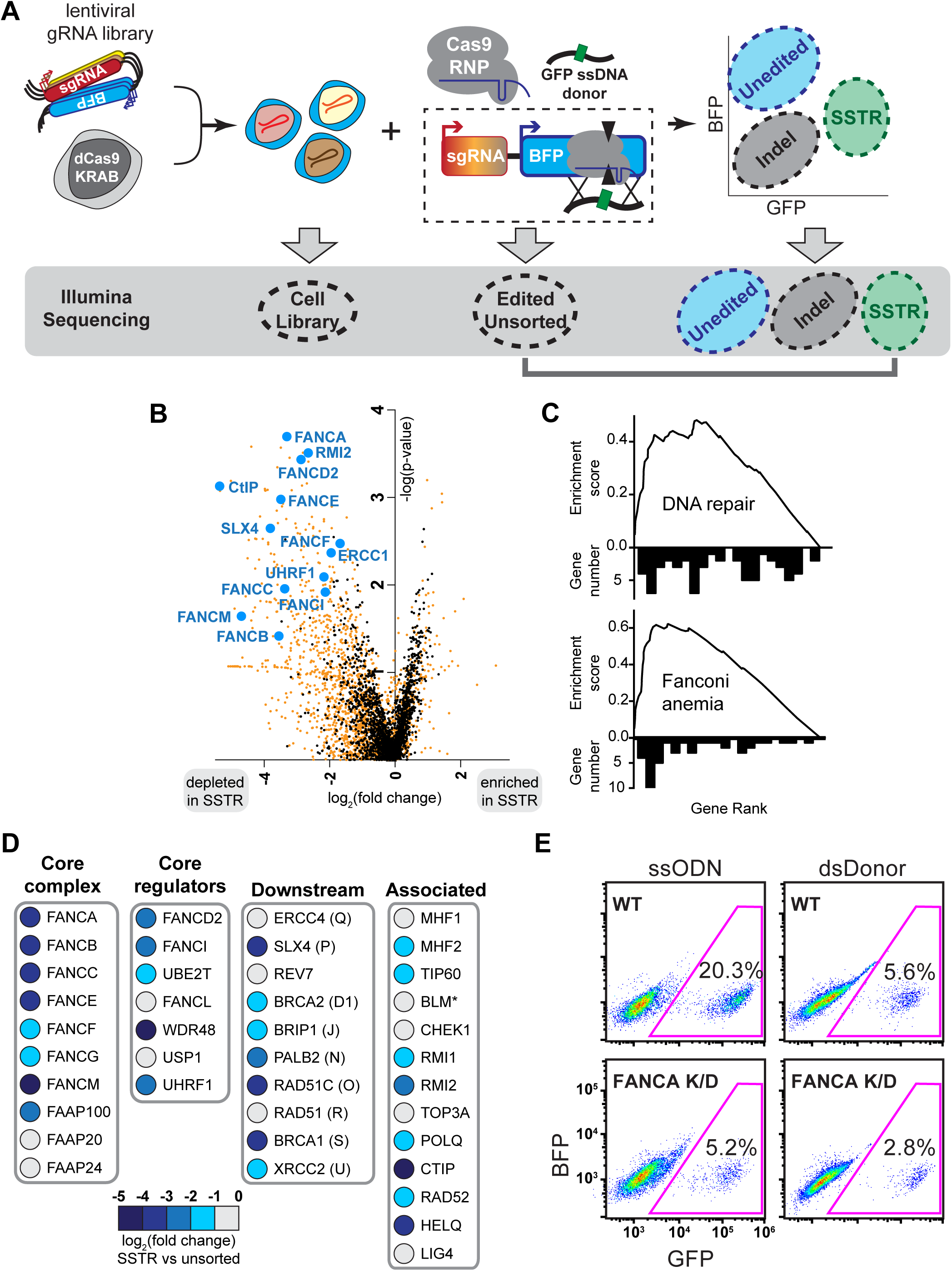
A coupled knockdown-editing screen identifies multiple Fanconia Anemia factors necessary for SSTR. **(A)** Schematic of the coupled knockdown-editing screening strategy. Lentiviruses comprising a BFP reporter and a gRNA are pooled and transduced into dCas9-KRAB K562 cells to produce a knockdown cell library. The gRNA library used here targets 2,000 distinct genes with gene ontology terms related to DNA (10,000 total gRNAs, 5 gRNAs per gene). This knockdown cell library is nucleofected with a pre-formed Cas9 RNP targeting BFP and an ssODN programming a 3bp mutation that converts BFP to GFP. Edited unsorted cells can be separated into three populations by FACS: Unedited (BFP+/GFP-); Indel (BFP-/GFP) and SSTR (BFP-/GFP+). Genes involved in regulating SSTR are identified by measuring guides that are significantly enriched or depleted in the SSTR population relative to the unsorted population by Illumina sequencing. **(B)** Multiple genes in the Fanconi Anemia pathway are depleted from the SSTR population. Representative SSTR genes are highlighted in blue, negative control untargeted gRNAs are shown in black, and targeted gRNAs are shown in orange. **(C)** GSEA analysis reveals that genes depleted from the SSTR population are significantly enriched in the FA repair pathway. **(D)** Multiple components of the FA repair pathway play a role in SSTR. The FA pathway can be separated into four functional categories: the FA Core complex, Core regulator factors involved in FANCD2-FANCI ubiquitination, Downstream repair effectors, and Associated factors involved in regulating FA outcomes. Genes are colored by log_2_ depletion of the indicated genes from each complex. Asterisk denotes gene scores that have been adjusted based on secondary analyses. Raw data is available in **[Document S3]**. **(E)** Individual CRISPRi knockdown of FANCA prevents SSTR. Representative BFP-GFP flow cytometry profiles are shown for untargeted (WT) or FANCA CRISPRi edited by co-administration of RNP targeting the BFP locus and ssODN or dsDonor. qPCR verifying knockdown of genes can be found in [**Extended Data Figure 6**].

To enable discovery of relatively low frequency events, we created a focused CRISPRi lentiviral library containing 2,000 genes (10,000 gRNAs, 5 gRNAs targeting each primary gene transcript) with gene ontology terms related to DNA processing **[Document S2 GUIDES]**^11^. This library was stably transduced at low multiplicity of infection into cells expressing dCas9-KRAB and selected for the gRNA construct for ten days to allow gRNA populations to reach equilibrium and for gRNAs targeting essential genes to drop out of the population. We harvested a sample of cells at this point as a control for comparison with previously published essentiality screens. We then electroporated cells with Cas9 RNP and the BFP-to-GFP ssODN^7^. Under unperturbed conditions, this combination of reporter, RNP, and ssODN yields ∼70% gene disruption (no longer BFP+) and ∼20% SSTR (BFP edited to GFP) [**Extended Data Figure 2**]. We harvested another sample of cells seven days after electroporation to identify genes whose knockdown is synthetic lethal with a Cas9-induced DSB, as measured by depletion only after introduction of Cas9. To identify genes involved in editing events, we used FACS to separate cells into unedited (BFP+/GFP-), Indel (BFP-/GFP-) and SSTR edited (BFP-/GFP+) populations **[Figure 1A, Extended Data Figure 2].** We used Illumina sequencing to measure gRNA abundances in each population, and compared these distributions to the edited unsorted cell population to reveal which target genes promote (gRNA depleted from edited population) or restrict (gRNA enriched from edited population) specific genome editing activities.

To benchmark our screening system, we identified essential genes by comparing the library-infected dCas9-KRAB CRISPRi cells with cells infected with only the gRNA library and no dCas9-KRAB **[Extended Data Figure 3A, Document S3 Essential Genes]**. Genes that were depleted after 14 days from the functional CRISPRi cells as compared to the gRNA-only control cells were significantly enriched for critical biological processes (DAVID^12^ analysis: proteasome core complex p=8.6e-14 and DNA replication p=1.7e-11). Furthermore, genes we identified as essential reproduced previously published essentiality screens **[Extended Data Figure 4A]**^11^, demonstrating that we had achieved stable gene knockdown and robust hit calling from the cell pools.

We next investigated genes whose knockdown was synthetic lethal with a Cas9-induced DSB. gRNAs targeting these genes should be depleted after Cas9 editing as compared to unedited cells **[Extended Data Figure 3B]**. While essential genes are progressively lost from the cell population over time **[Extended Data Figure 4B]**, genes whose knockdown is synthetic lethal with DNA damage are lost only after a Cas9 DSB **[Extended Data Figure 4C]**. Genes required to survive a single Cas9-induced DSB were enriched for the GO terms such as cell cycle arrest (p=3.3E-05) and response to DNA damage (p= 3.4E-22) and include several factors previously reported to be synthetic lethal with other DNA damaging agents **[Document S3 SurviveDSB]**. For example, knockdown of MYBBP1A has recently been reported to cause senescence in combination with nonspecific DNA damage^13^. Our screening cell line contains a single copy of the targeted BFP allele, which suggests that a single DSB is sufficient to trigger genotoxic-induced senescence when these synthetic lethal genes are depleted. Together, our results indicate that our coupled inhibition-editing strategy performs well in identifying not only essential genes, but also genes involved in DNA repair pathways required to survive a Cas9 DSB. Future investigation of novel genes identified as synthetic lethal with a DSB could provide new insight into mechanisms of genome surveillance.

To identify factors required for SSTR editing, we used FACS to isolate GFP+ cells (that had undergone BFP-to-GFP conversion via SSTR) and measured depletion of gRNAs relative to the unsorted edited pool **[Figure 1B]**. Strikingly, 70% (28/40) of genes annotated in the Fanconi Anemia (FA) pathway were robustly and consistently depleted from the GFP+ population relative to unsorted edited cells. Gene set enrichment analysis^14^ verified that DNA repair in general and the FA pathway in particular was a defining feature of SSTR **[Figure 1C]**. Several distinct functional groups within the FA repair pathway were identified as required for SSTR: multiple components of the FA core complex that senses lesions, FA core regulatory components that activate the FANCD2-FANCI heterodimer, downstream effector proteins that repair lesions, and associated proteins that interact with canonical FA repair factors **[Figure 1D]**. Our identification of the FA pathway as central to SSTR was striking, as to the best of our knowledge the FA pathway has not previously been investigated for its role in Cas9 gene editing.

We used individual knockdown of FANCA, RAD51, and other DNA repair genes to further investigate the genetic basis of SSTR and dsDonor HDR. Previous reports have indicated the editing outcomes of SSTR are RAD51-independent and ineffective during G2/M^15,16^, while dsDonor HDR is RAD51-dependent^17^, FANCA-dependent^18^, and active during G2/M^19^. Using Cas9 RNPs to edit the same locus with dsDonors and ssODNs in K562 human erythroleukemia cells, we found approximately four-fold higher gene replacement efficiency with ssODNs. Knockdown of FANCA caused statistically significant four-fold reduction in SSTR (p<0.05, Welch’s two-sided t-test) and a non-significant two-fold reduction in dsDonor HDR (p=0.22, Welch’s two-sided t-test) **[Figure 1E]**. These results highlight the unexpected role of FANCA in SSTR and suggest it might also play some role in HDR. As expected, neither SSTR nor HDR required NHEJ (mediated by LIG4), or the related Alternative End Joining (Alt-EJ) pathway (mediated by PARP1). Notably, we found that knockdown of RAD51 abolished dsDonor HDR but had no effect upon SSTR **[Extended Data Figure 5]**. Taken together, individual knockdown bolsters the hypothesis derived from the primary screen that SSTR is a genetically distinct pathway from dsDonor-mediated HDR.

The FA repair pathway is best understood in its capacity to identify and repair interstrand crosslinks (ICLs) throughout the genome, but has recently gained attention for its role in protecting stalled replication forks^20^-^22^. In the presence of ICLs or a stalled fork, the FA core complex (comprised of FANCA, B, C, E, F, G, L, M, FAAP100, FAAP20, and FAAP24) is required for monoubiquitination and activation of the FANCD2-FANCI heterodimer by UBE2T and FANCL. Monoubiquitination then leads to recruitment of downstream factors that repair the lesion via nucleotide excision repair (NER) or specialized homologous recombination sub-pathways. Subsequent to repair, FANCD2/FANCI is recycled through deubiquitination by USP1 and WDR48. Deactivation of FANCD2/FANCI appears to be a key step in restoring homeostasis, as mutants in USP1 and WDR48 phenocopy classical FA mutants with an increased sensitivity to ICL-causing agents. Notably, our screen identified that SSTR depends upon genes that act in every functional category of the FA repair pathway **[Figure 1D].** SSTR is therefore likely to be a central activity of the FA repair pathway as opposed to a moonlighting activity of one or more FA genes.

To further explore the genetic basis of SSTR, we used CRISPRi to stably knock down seven separate FA repair genes that operate at different places in the FA pathway and quantified the frequency of Cas9-mediated SSTR at multiple loci. Knockdown of FANCA, FANCD2, FANCE, FANCF, FANCL, HELQ, UBE2T, USP1, and WDR48 all substantially decreased SSTR at a stably integrated BFP reporter, as measured by flow cytometry **[Figure 2A]**. Stable cDNA re-expression of each factor restored wildtype levels of SSTR, demonstrating that CRISPRi was specific to the targeted gene and that ablation of each gene was solely responsible for the loss of SSTR. Re-expression of an FA factor in the context of its knockdown increased editing efficiency up to 8-fold. These results demonstrate that multiple genes in different parts of the FA repair pathway are required for SSTR editing, that their presence is necessary for efficient SSTR, and that re-expression is sufficient to restore SSTR.

**Figure 2:**
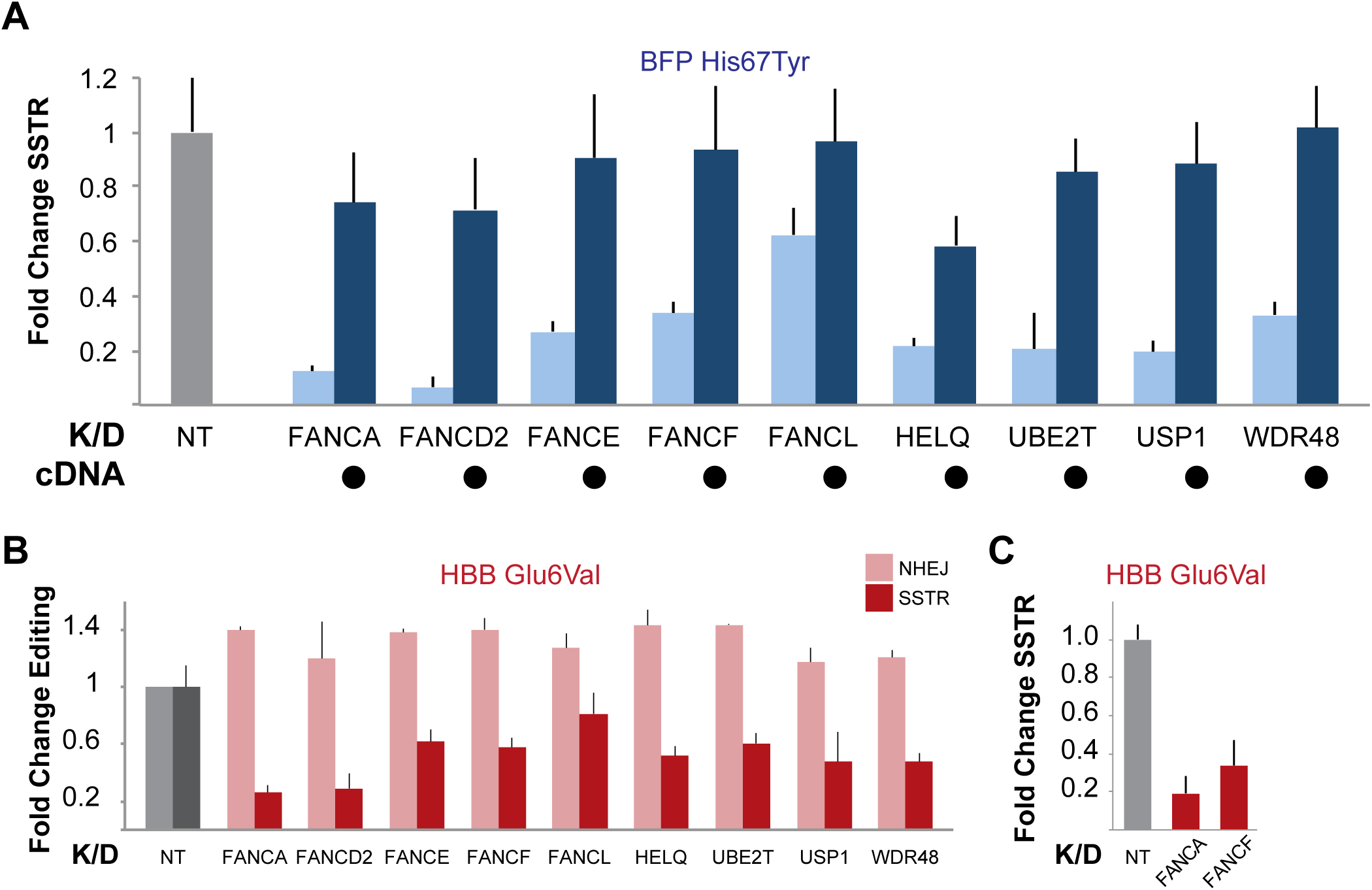
Knockdown of FA genes specifically inhibits SSTR. **(A)** Knockdown of multiple FA repair genes prevents SSTR at a single copy genomically integrated BFP reporter. Stable cDNA re-expression rescues this phenotype. Stable CRISPRi cell lines with untargeted (NT, grey) gRNA or gRNA targeting specific genes were edited by co-administration of RNP targeting the BFP reporter and an ssODN containing a BFP→GFP mutation (see **[Extended Data Figure 6]** for transcript levels). The frequency of GFP+ cells was quantified by flow cytometry, normalized to untargeted controls, and %SSTR was plotted for knockdown of FA factors (light blue) or knockdown of FA factors in the context of cDNA re-expression (dark blue). Data are presented as mean±sd of at least two biological replicates. **(B)** Knockdown of FA repair genes prevents SSTR and upregulates NHEJ at the HBB Glu6 codon. Cell lines with untargeted control gRNA (NT, grey), or gRNAs targeting each FA gene (red) were edited using Cas9 targeting the HBB locus and a ssDNA donor that introduces the Glu6Val sickle mutation. NHEJ (light columns) and SSTR (dark columns) were measured by amplicon Illumina sequencing of edited alleles **[Extended Data Figure 8]** and normalized to untargeted controls. Data presented are the mean±sd of at least two biological replicates. **(C)** Inhibition of FA repair genes prevents SSTR in primary fibroblasts. Human neonatal dermal fibroblasts were treated without (WT) or with siRNA targeting the indicated genes and edited as described in **[Figure 2B]**. Verification of knockdown can be found in **[Extended Data Figure 5C]**.

In addition to the FA pathway’s well-characterized roles in ICL repair, there is an emerging view that it plays additional roles in preserving genome stability. FA genes protect against aberrant chromosomal structures and replication stress via specialized subcomplexes that in part depend upon particular helicases, including Bloom’s helicase (BLM) and the 3’-5’ ssDNA helicase HELQ. We found that siRNA knockdown of BLM had no effect on SSTR **[Extended Data Figure 7],** but knockdown of HELQ markedly reduced SSTR **[Fig 2A**, **Extended Data Figure 7]**. BLM and its interaction partner RMI2 exhibited strong phenotypes in the primary screen **[Document S3, SSTR]**. However, both of these factors were required (p<0.01) for survival of a Cas9-induced DSB **[Document S3, SurviveDSB]**, which suggests a role for the BLM complex in surviving a DSB instead of SSTR itself. While BLM has been linked to FA-mediated resolution of replication stress^23^, HELQ directly interacts with FANCD2/FANCI with unknown functional significance^24^. HELQ also interacts with multiple recombination subcomplexes, including BCDX2 (RAD51B, RAD51C, RAD5D, and XRCC2) and CX3 (RAD51C-XRCC3). These complexes could promote recombination between the ssODN and genomic DNA, and we asked if these complexes also impact SSTR.

We found that RAD51C is required for SSTR, but RAD51B and XRCC2 are not. This suggests that BCDX2 does not play a role in SSTR **[Extended Data Figure 7]**. Conversely, both RAD51C and XRCC3 are required for SSTR, implicating the CX3 complex in SSTR [**Extended Data Figure 7**]. Intriguingly, CX3 has been reported to act downstream of RAD51 filament formation^25^, but we found that RAD51 itself is dispensable for SSTR [**Figure 1B, Extended Data Figure 5**]. We anticipate that future work to characterize how the FA pathway interacts with downstream effectors, especially polymerases and genes that mediate recombination, will provide valuable insights into the mechanism of SSTR and its interaction with other pathways that maintain genome stability.

Inhibition of SSTR by interfering with the FA pathway could work by globally reconfiguring DNA repair pathway preference or by specifically inhibiting SSTR. We investigated how the FA pathway influences repair pathway choice by inhibiting several FA genes and measuring editing outcomes using Illumina sequencing **[Extended Data Figure 8]**. When editing the endogenous hemoglobin β (HBB) locus at the causative amino acid (Glu6) for sickle cell disease, we found that all seven FA factors are required for SSTR editing **[Figure 2B]**. Notably, knockdown of FA genes decreased levels of SSTR while simultaneously increasing levels of NHEJ, such that total editing (SSTR + gene disruption) remained relatively constant **[Figure 2B, Extended Data Figure 9**]. However, when we edited the HBB locus in the absence of an ssODN, we found that knockdown of FA repair genes did not significantly increase NHEJ frequency on its own **[Extended Data Figure 9]**. We found similar results at the BFP locus when measuring editing outcomes by Illumina sequencing. These results imply that the FA repair pathway acts to divert repair events that would otherwise be repaired by NHEJ into SSTR outcomes. This model parallels proposed roles for the FA pathway in balancing NHEJ and HR repair frequencies during ICL repair^26,27^ and balancing Alt-EJ, NHEJ, and HDR repair outcomes near DSBs^28^.

To determine if FA repair genes are responsible for SSTR in primary human cells, we edited human neonatal dermal fibroblasts at HBB Glu6. These fibroblasts have previously been shown to be capable of SSTR repair, albeit at lower levels than many cell lines^15^. Untreated or mock siRNA treated fibroblasts exhibited approximately 5% SSTR at the HBB locus, as measured by Illumina sequencing. siRNA knockdown of either FANCA or FANCE led to an approximately five-fold reduction in SSTR **[Figure 2C]**. Therefore, the FA repair pathway is tightly linked to SSTR in at least one primary human cell type.

The sequence outcomes of genomic disruption (indels) following Cas9-induced DSBs are often nonrandom and surprisingly consistent at individual loci, leading to an emerging model that repair outcomes are determined by the intrinsic repair pathway preferences of the edited cell and the sequence immediately adjacent to the cut site^29^. To determine how FA pathway disruption affects the characteristic spectrum of indels as a Cas9-induced break, we characterized individual allele frequencies in unperturbed and FA knockdown CRISPRi cell lines using Illumina sequencing at both BFP and HBB Glu6. We also examined SSTR conversion tracts, a function of SNP integration relative to distance from the Cas9-induced break, by following the incorporation of several single nucleotide polymorphisms (SNPs) encoded by the ssODN into the genomic sequence.

In the absence of an ssODN to program SSTR, neither the overall frequency nor the pattern of indels at the Cas9 cut site was affected by disruption of the FA repair pathway **[Figure 3A]**. We furthermore observed no change in indel spectra upon FA knockdown when editing cells in the presence of an SSTR-templating ssODN. However, when editing with an ssODN, SSTR decreased dramatically upon disruption of the FA pathway. This decrease was remarkably uniform across SNPs within the ssODN and did not measurably alter the SNP conversion tracts. These results reinforce our earlier observation that FA repair pathway inactivation specifically inhibits SSTR without altering the frequency of indels. Additionally, the molecular sequence outcomes of NHEJ are unaffected by the FA pathway. Instead, the FA pathway is restricted to SSTR repair and the balance between NHEJ and SSTR, but does not play a direct role in error-prone end-joining pathways.

**Figure 3:**
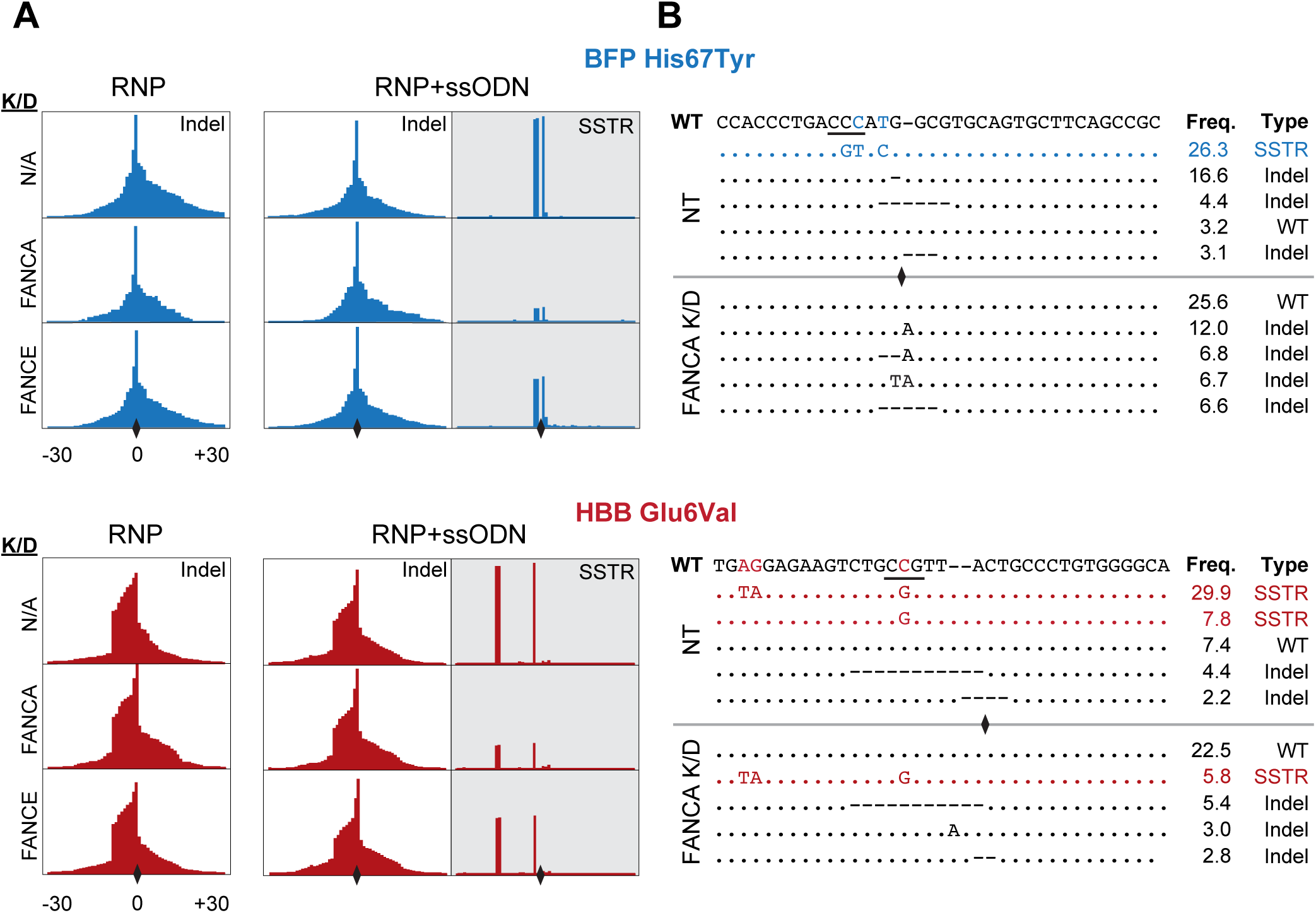
FA gene knockdown rebalances edited alleles without affecting the spectra of molecular outcomes. **(A)** FA repair genes are required for SSTR but not end-joining. Cell lines with stable knockdown of control (untargeted gRNA, NT), FANCA, or FANCE were edited at the BFP (blue) or HBB Glu6 (red) loci. Editing was performed without (RNP only, left panels) or with (RNP+ssODN, right panels) ssODN containing three trackable SNPs and editing outcomes were measured using amplicon sequencing. The frequency of sequence alteration by insertion/deletion (indel) or SSTR is plotted as a function of distance from the Cas9 cut site (diamond). Frequency of sequence alteration is displayed from 30bp PAM proximal (-) to 30bp PAM distal (+) of the Cas9 cut site. Data displayed are representative of biological replicates. **(B)** SSTR is the most common allele in Cas9-edited cell populations and is lost when FA genes are knocked down. Allele frequencies were calculated for the editing experiments presented in **[Figure 3A]**. Wildtype sequences with PAM underlined are presented for BFP and HBB Glu6 along with the sequence alignments of the five most common alleles produced when editing the BFP or HBB loci in the context of control (NT) or FANCA knockdown. Alleles are categorized as SSTR, wildtype (WT), or Indel. Percentages of total reads are presented for each allele.

In sum, we have found that multiple functional complexes within the FA repair pathway are necessary for Cas9-mediated SSTR. Genome editing is commonly grouped into two categories, genetic disruption and genetic replacement, based on sequence outcomes^2^. Our results demonstrate that final genetic replacement outcomes using different templates (ssODN vs dsDonors) are identical at the sequence level but stem from completely different pathways **[Figure 4]**. Specifically, information from double stranded DNA templates and genomic DNA are incorporated using Rad51-dependent processes, but single stranded DNA templates are incorporated through the FA pathway. A great deal of work has focused on improving HDR during gene editing by activating Rad51-mediated processes, including Rad51 agonist small molecules^30^ and strategies to stimulate recruitment of Rad51 throughout the cell cyle^31^. Our results indicate that future efforts to the activate FA pathway could be invaluable during gene editing for research or therapeutic uses.

**Figure 4:**
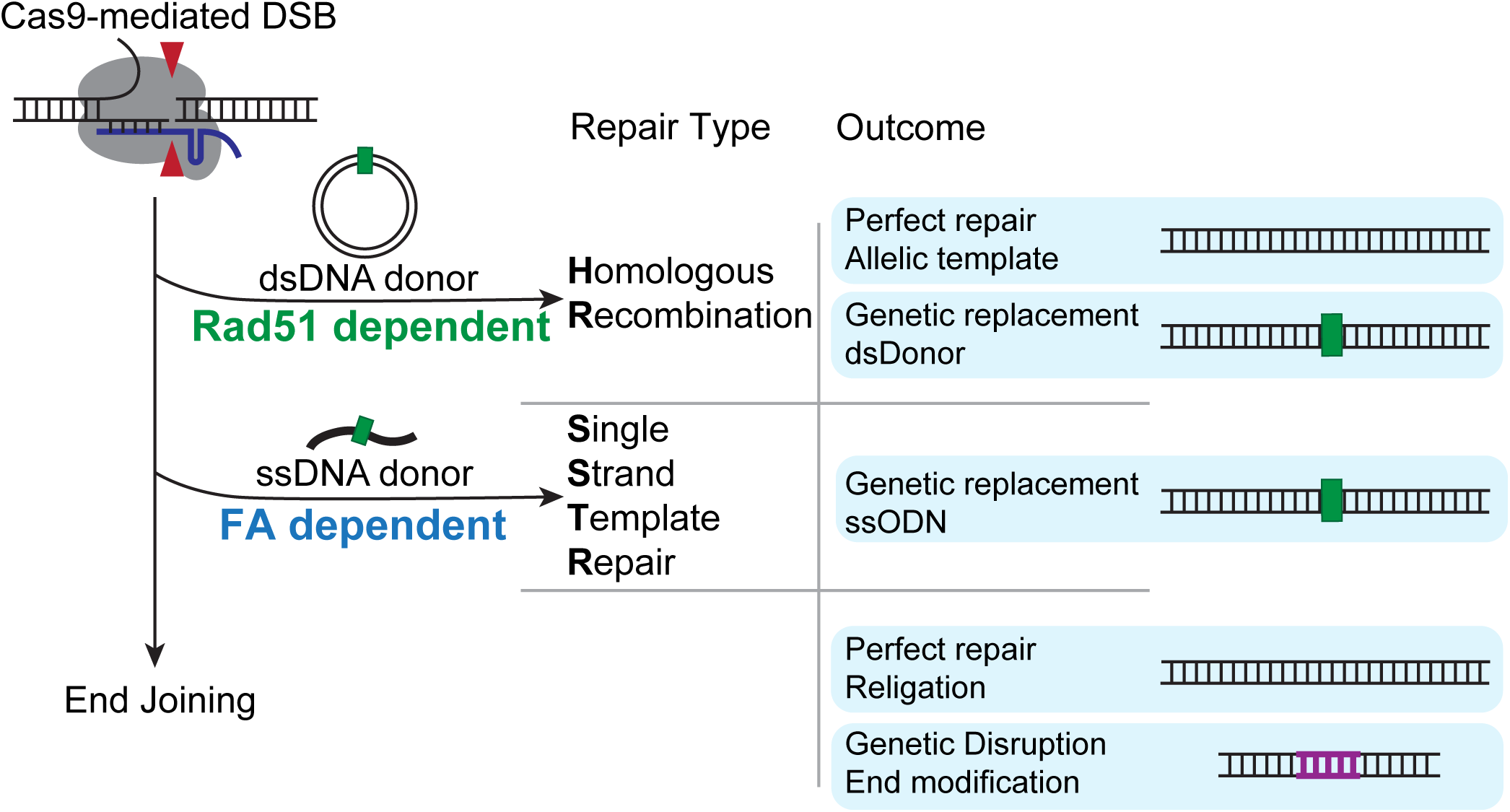
Cas9-mediated genomic replacement can be separated into two distinct pathways based on the type of template used for repair. Cas9-induced DSBs can be repaired using donor-dependent or end joining repair events. Donor-dependent events can be separated into Rad51-dependent FANCA-independent (HR) or Rad51-independent FA-dependent (SSTR) repair. Sequence replacement outcomes are in competition with end-joining, whose outcomes include perfect repair of the original sequence, genetic replacement, and genetic disruption.

Cas9-mediated genome editing holds great promise for the treatment of genetic diseases such as sickle cell disease and Fanconi Anemia. High rates of gene editing are typically required for therapeutic editing applications, but editing efficiencies can differ greatly between cells. Without knowledge of the pathways responsible for genetic replacement outcomes and the activity of those pathways in the targeted cell type, it was previously difficult to rationalize why editing might fail in one application while succeeding in another. Our results predict that human cell types with intrinsic repair preferences that impact the FA pathway will be more or less capable of SSTR **[Extended Data Figure 1].** The expression level of FA-related factors could in future be useful as a biomarker for patient cell “editability”, and treatments that enhance the activity of the FA pathway could be especially valuable in difficult to edit cells. For example, we found that complementing FA pathway knockdown yields up to 8-fold increase in editing efficiency in cell lines **[Figure 2A]**. This suggests that reactivating the FA pathway could be valuable in cases where it has been disrupted, such as in Fanconi Anemia itself. Small molecule activators of the FA pathway remain to be identified, but our results suggest that transiently increasing the levels of FA proteins could complement patient-specific defects to enable lasting gene editing cures. More broadly, our results suggest that patient genotype or transcriptome could increase or decrease the effectiveness of therapeutic treatments in previously unanticipated ways. Deeper understanding of the molecular basis of SSTR and dsDonor HDR is likely to suggest new biomarkers to ‘match’ patient genotype with therapeutic editing strategy.

Finally, our data imply that the default repair pathway for DSBs, especially DSBs introduced by Cas9, is end joining, and that the activity of the FA repair pathway determines whether many events will instead be repaired by SSTR **[Figure 2A, Figure 4]**. Cas9 is very stable on genomic targets^7,32^, and so it is possible that Cas9 itself is recognized as an interstrand crosslink or roadblock within the genome. However, we disfavor this hypothesis because FA knockdown only impacts SSTR without directly affecting indels. Instead, we hypothesize that Cas9-stimulated repair using an ssODN template mimics some substrate of the FA pathway, such as a stalled replication fork. We note that SSTR is much more efficient than HDR from a double stranded DNA template **[Figure 1E]**, to the extent that in many cell lines, the most common single allele at an edited locus is the product of SSTR **[Figure 3B]**. This ability raises intriguing questions about genome integrity in the presence of single stranded DNA exposed by R-loops, replication crises, or viral infection. The FA pathway has already been implicated in replication crises, and future work to address remaining questions could provide insight into mechanisms by which human cells maintain their genomes.

---

**Extended Data Figure 1:**
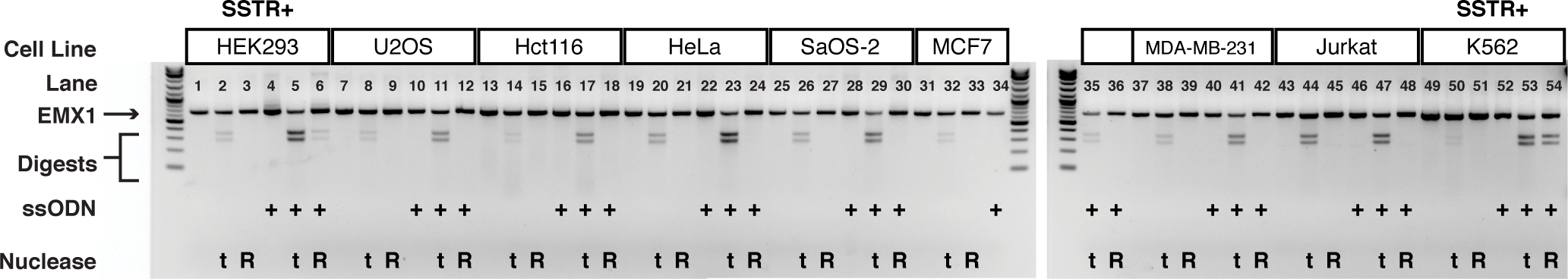
SSTR efficiency varies in different human cell lines. Nine cell lines were edited using RNP targeting the EMX1 locus either without or with ssODN containing PciI sequence. The edited locus was amplified, re-annealed, and digested using T7 Endonuclease I (t), which quantifies gene disruption, or the restriction enzyme PciI (R), which quantifies SSTR.

**Extended Data Figure 2:**
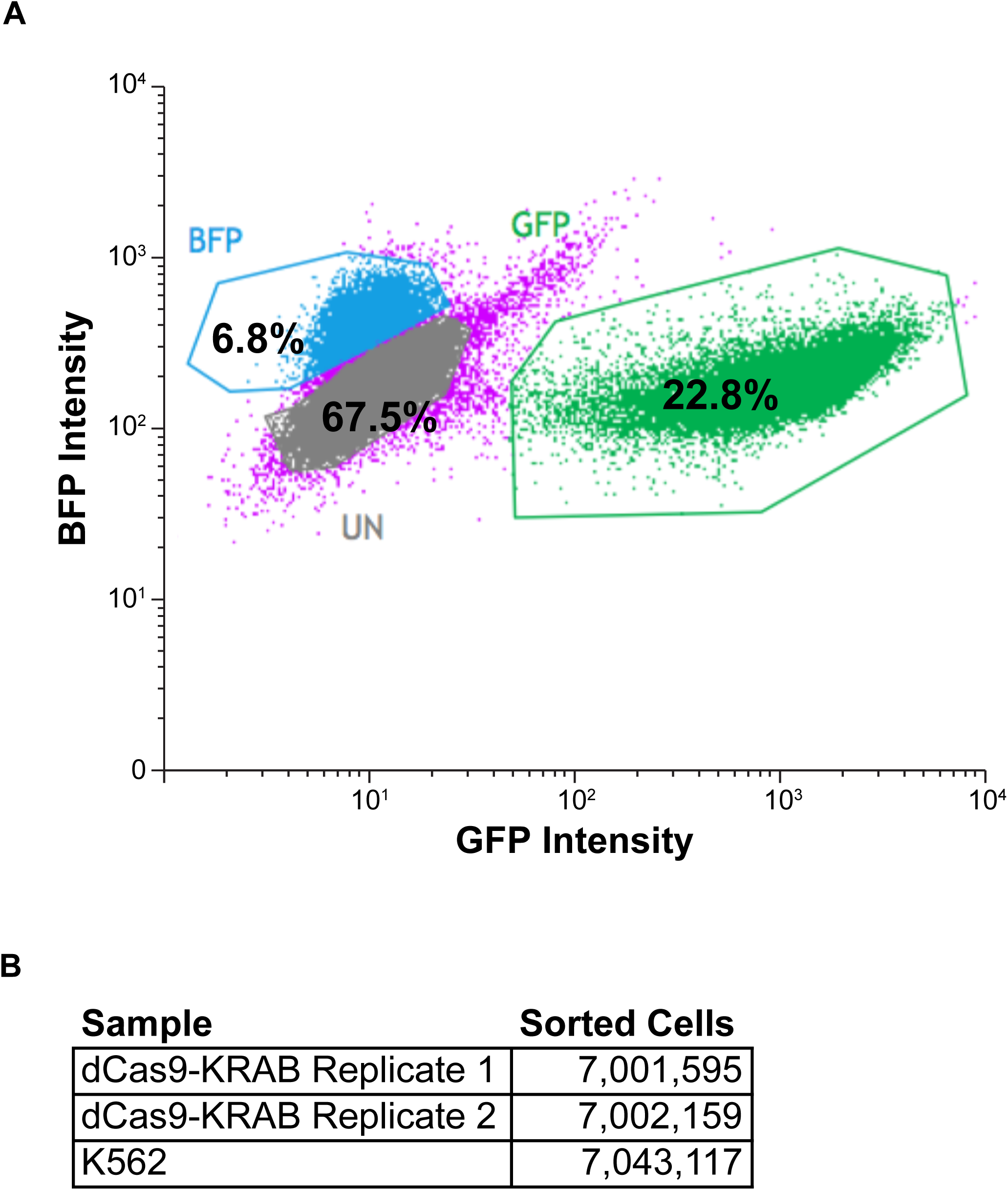
BFP, GFP, and Non-fluorescent populations are effectively resolved during a pooled CRISPRi screen. **(A)** Replicate one of dCas9-KRAB cells infected with gRNA library and edited at the BFP locus. Live single cells were plotted to compare BFP and GFP intensities. Three populations were sorted: BFP+/GFP-(BFP), BFP-/GFP+ (GFP), and BFP-/GFP-(UN). Percentages in each gate are presented for 100,000 cells. **(B)** Total cells sorted for each edited population.

**Extended Data Figure 3:**
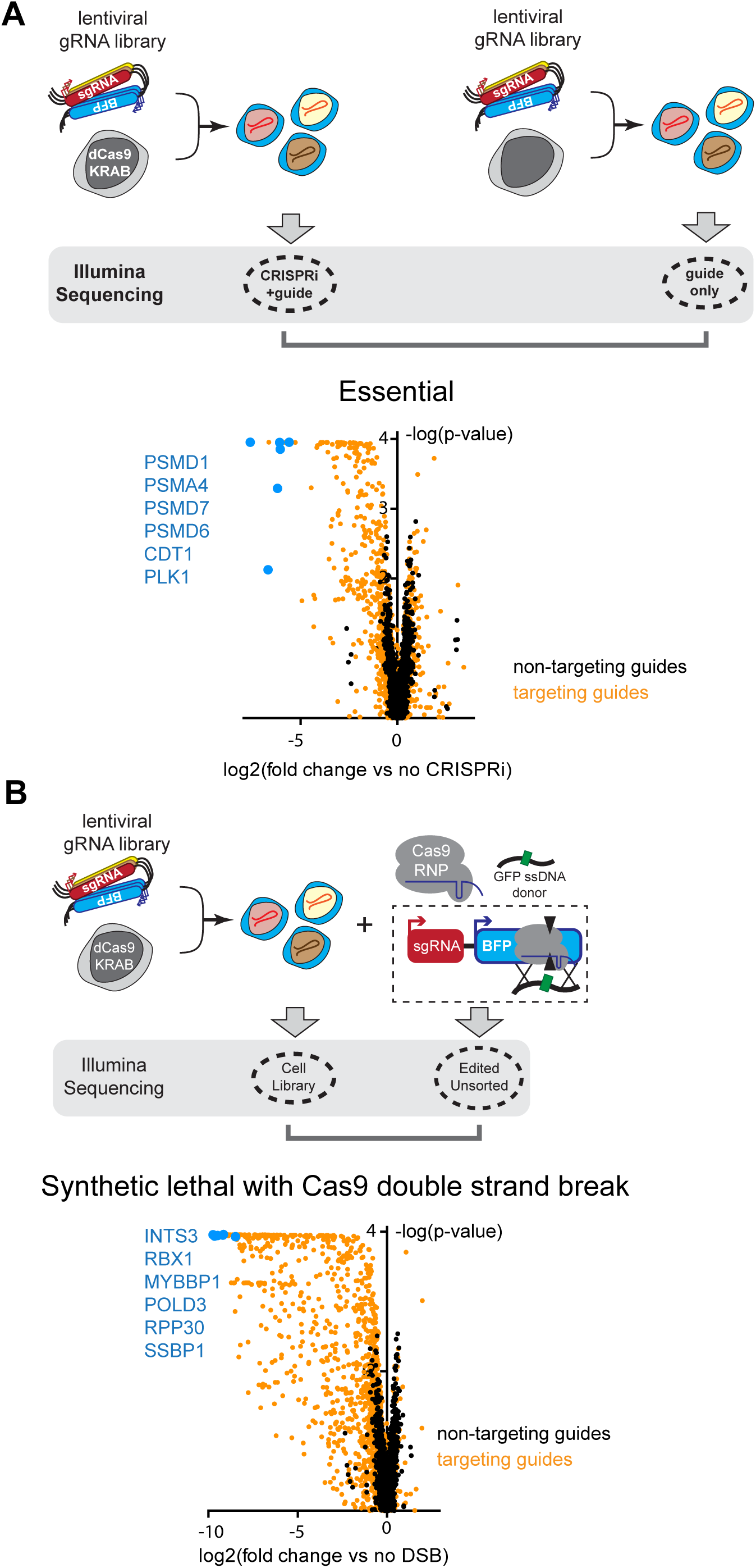
Pooled CRISPRi screens identify DNA repair genes that contribute to multiple phenotypes. (A) Essential genes in the K562 cell background. A volcano plot identifies genes from the DNA repair library enriched or depleted in CRISPRi cells relative to gRNA-only controls. Representative essential genes are highlighted in blue, negative control untargeted gRNAs are shown in black, and targeted gRNAs are shown in orange. (B) Genes synthetic lethal with a DSB. A volcano plot identifies genes from the DNA repair library enriched or depleted in CRISPRi cells treated with Cas9 relative to untreated cell populations. Representative synthetic lethal genes are highlighted in blue, negative control untargeted gRNAs are shown in black, and targeted gRNAs are shown in orange.

**Extended Data Figure 4:**
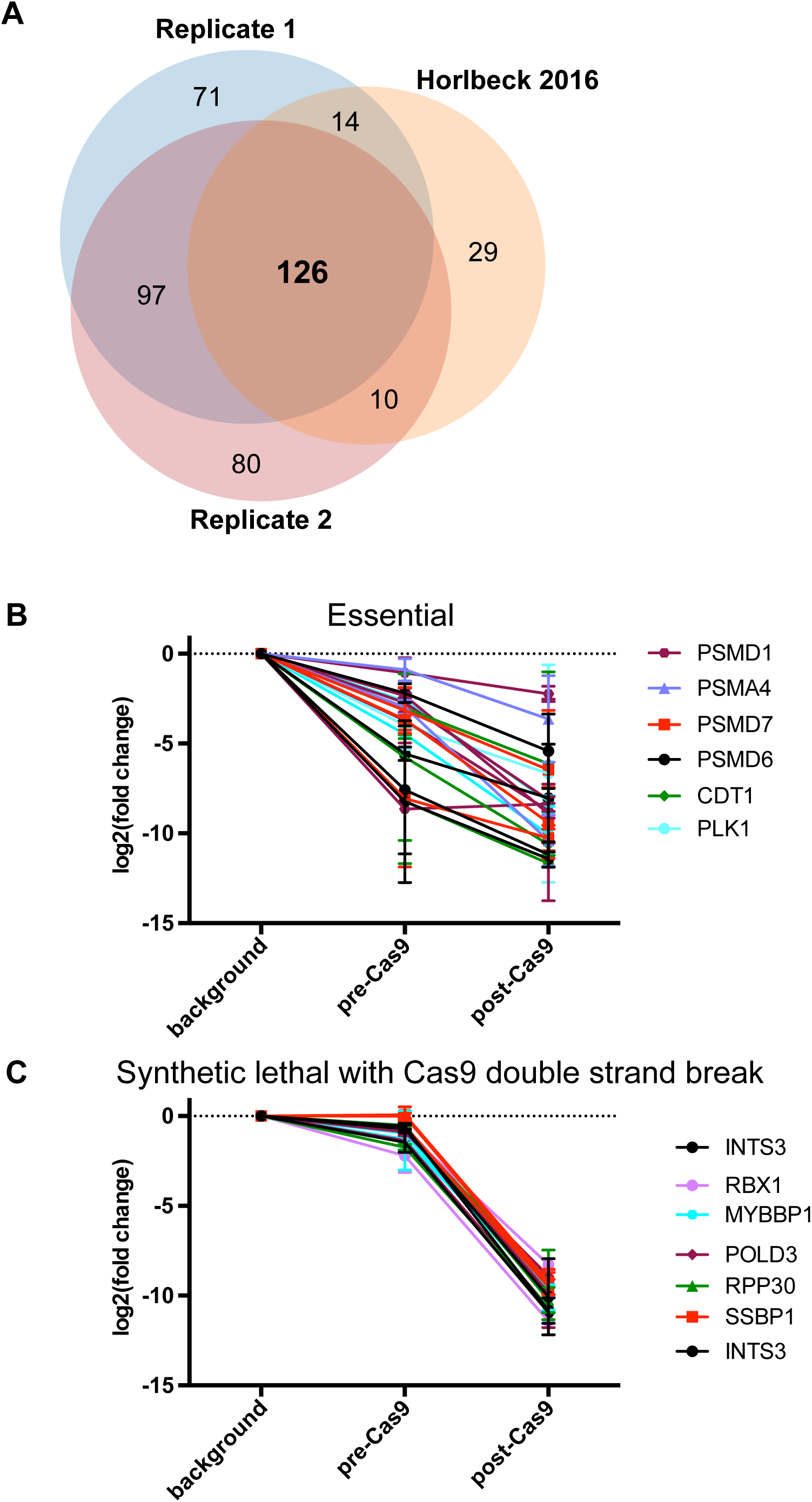
(A) Essential genes identified in this study substantially overlap with previous studies. The distribution of gRNAs in two unedited dCas9-KRAB cell libraries was separately compared to the gRNA distribution in an unedited K562 cell library and gene scores for each target genes were calculated. Essential genes were defined as those showing significant (p<0.05) depletion (log_2_(fold change)<-0.5) in the dCas9-KRAB population. These essential genes were compared to the K562 dataset generated in (Horlbeck et al, 2016), filtered for significance (p<0.05) and magnitude of effect (phenotype<-0.2). Overlap between gene lists is presented as a Venn diagram. (B) Essential genes are progressively lost from library. (C) Genes that are synthetically lethal with a single DSB are abruptly lost from library following Cas9 treatment.

**Extended Data Figure 5:**
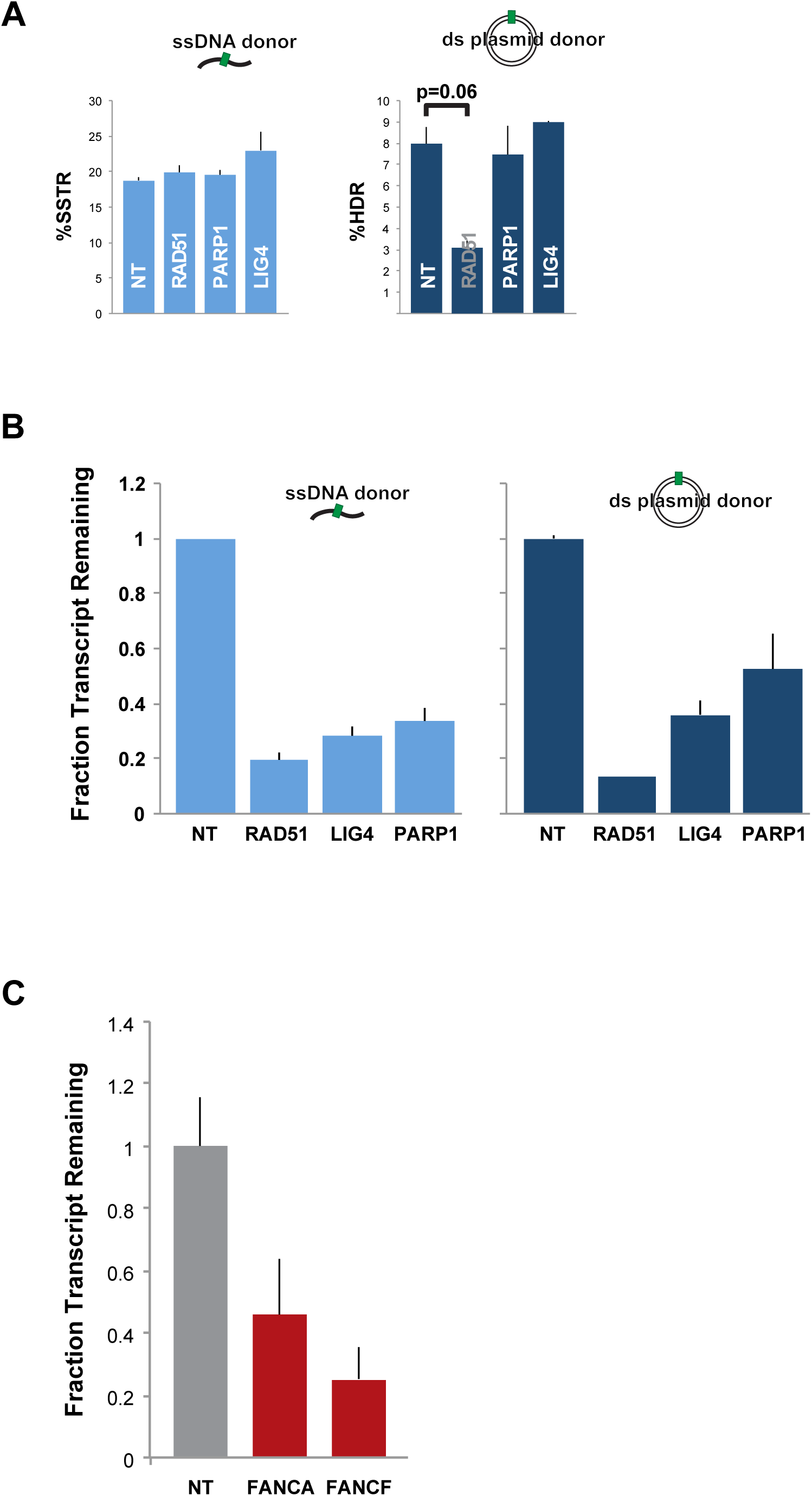
Effective knockdown of DNA repair factors using siRNA. **(A)** Knockdown and editing of DSB repair factors reveals genetic differences between HDR and SSTR. K562 cells expressing a BFP reporter were treated with the indicated siRNAs and edited by co-administration of RNP targeting the BFP reporter with ssODN or dsDonor containing a BFP->GFP mutation. **(B)** siRNA of RAD51, PARP1, and LIG4 in K562 cells **[Extended Data Figure 5A]**. Cells were siRNA treated for 48 hours prior to editing. Fold depletion of the siRNA-target transcript over controls (ACTB, GAPDH) was measured by qPCR. Data presented represent the transcriptional state of the cells at the time of editing. All data were calculated from technical triplicate and biological replicate. **(C)** siRNA knockdown of FANCA and FANCF in human dermal fibroblasts. Cells were siRNA treated prior to editing **[Figure 2C]** as described in **[Extended Data Figure 5B]**.

**Extended Data Figure 6:**
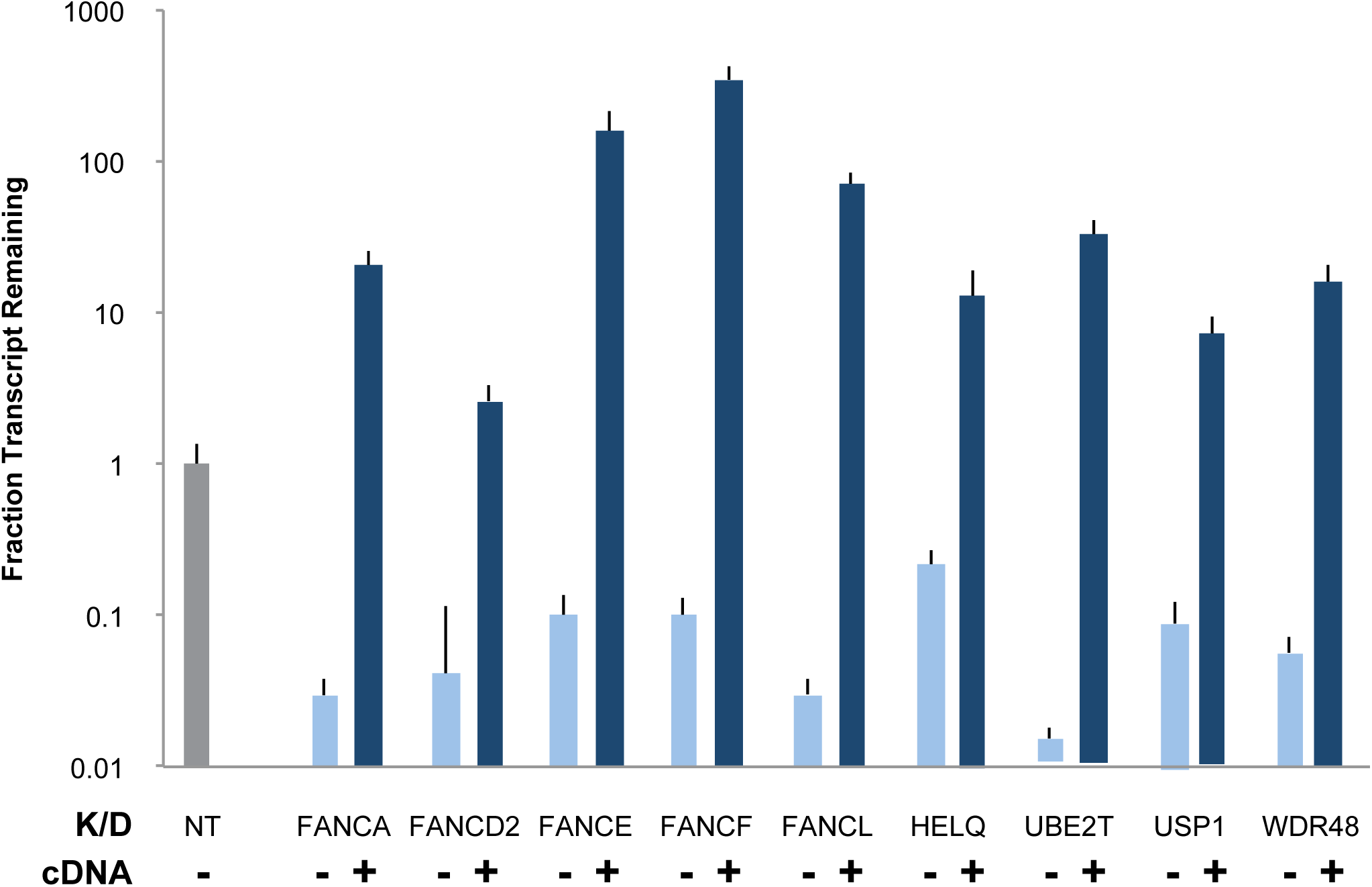
Effective knockdown of DNA repair factors using CRISPRi. **(A)** Stable CRISPRi cells targeting the indicated gene with or without re-expression of a cDNA of the indicated gene were harvested **[Figures 2A-B]** and fold depletion of the indicated transcripts over control transcripts (ACTB, GAPDH) was measured by qPCR. Data presented were calculated from technical triplicate and biological replicate.

**Extended Data Figure 7:**
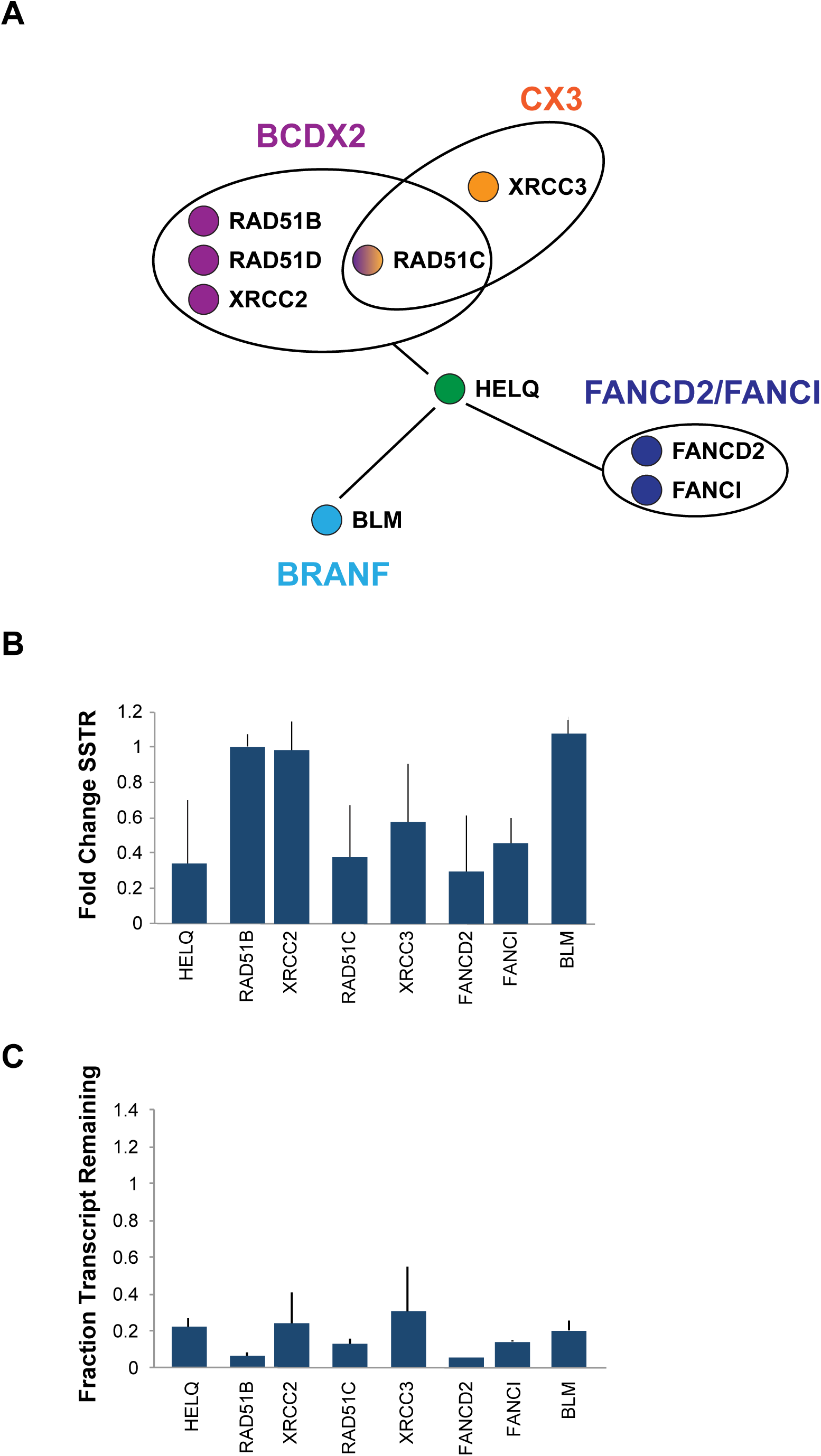
HELQ interaction partners play a role in SSTR. **(A)** Interaction map of HELQ reproduced from an earlier study (Adelman 2013) illustrates direct physical interactions between HELQ and other complexes and reported interactions from BIOGRID, STRING and MINT databases. **(B)** HELQ interaction partners play a role in SSTR. K562 cells were siRNA or CRISPRi treated against the indicated target genes prior to editing at the BFP locus. Data presented are the mean±sd of biological duplicate. **(C)** HELQ interaction partners can be effectively depleted by siRNA or CRISPRi. Fold depletion of the target transcript over controls (ACTB, GAPDH) was measured by qPCR. Data presented were calculated from technical triplicate and biological replicate.

**Extended Data Figure 8:**
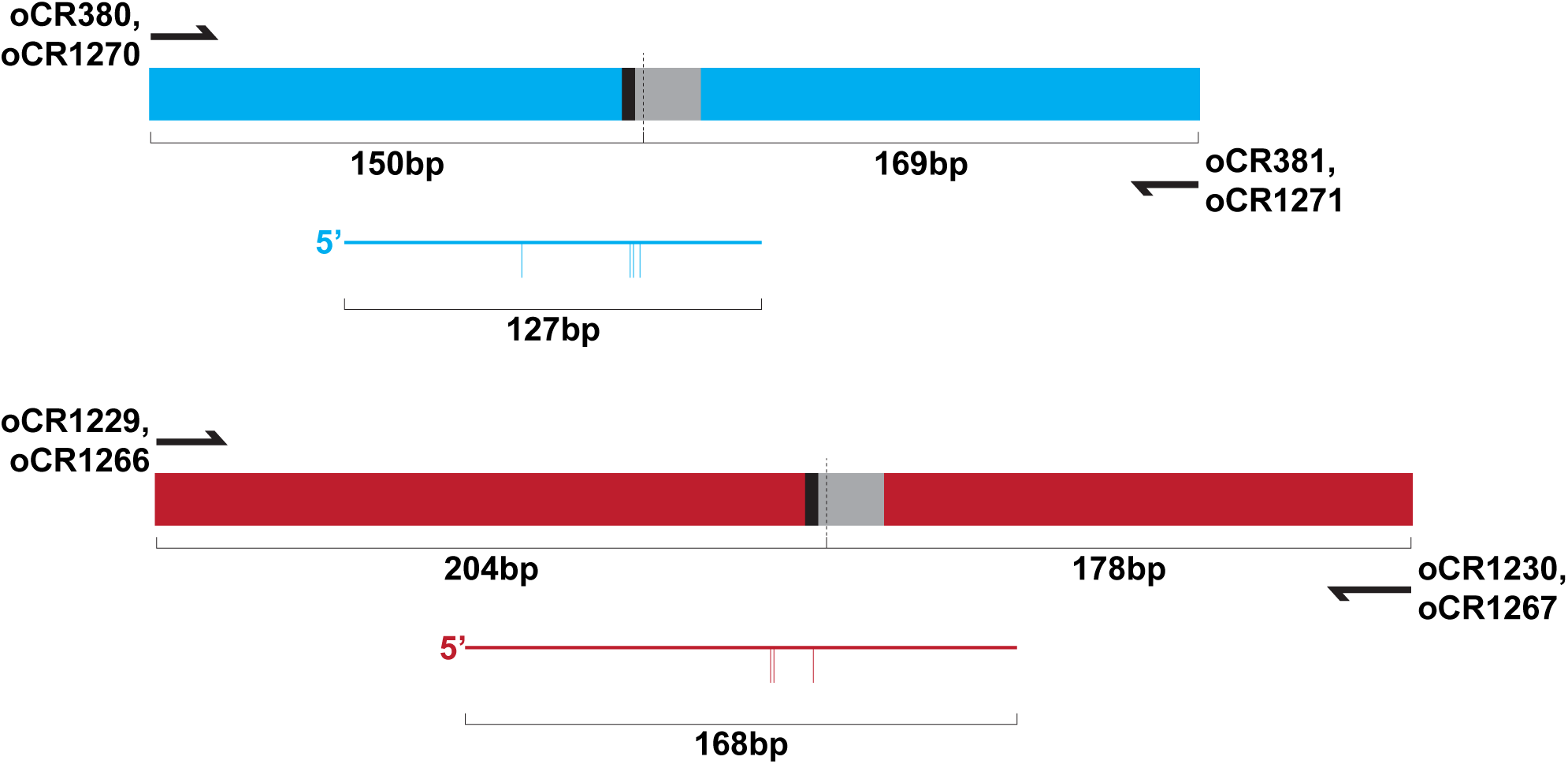
Schematic of amplicon sequencing used at BFP or HBB loci. Protospacer (grey) and PAM (black) are presented within amplified sequence. ssODN size and orientation is also displayed. Vertical hash marks indicate locations where ssODN and genomic sequences differ.

**Extended Data Figure 9:**
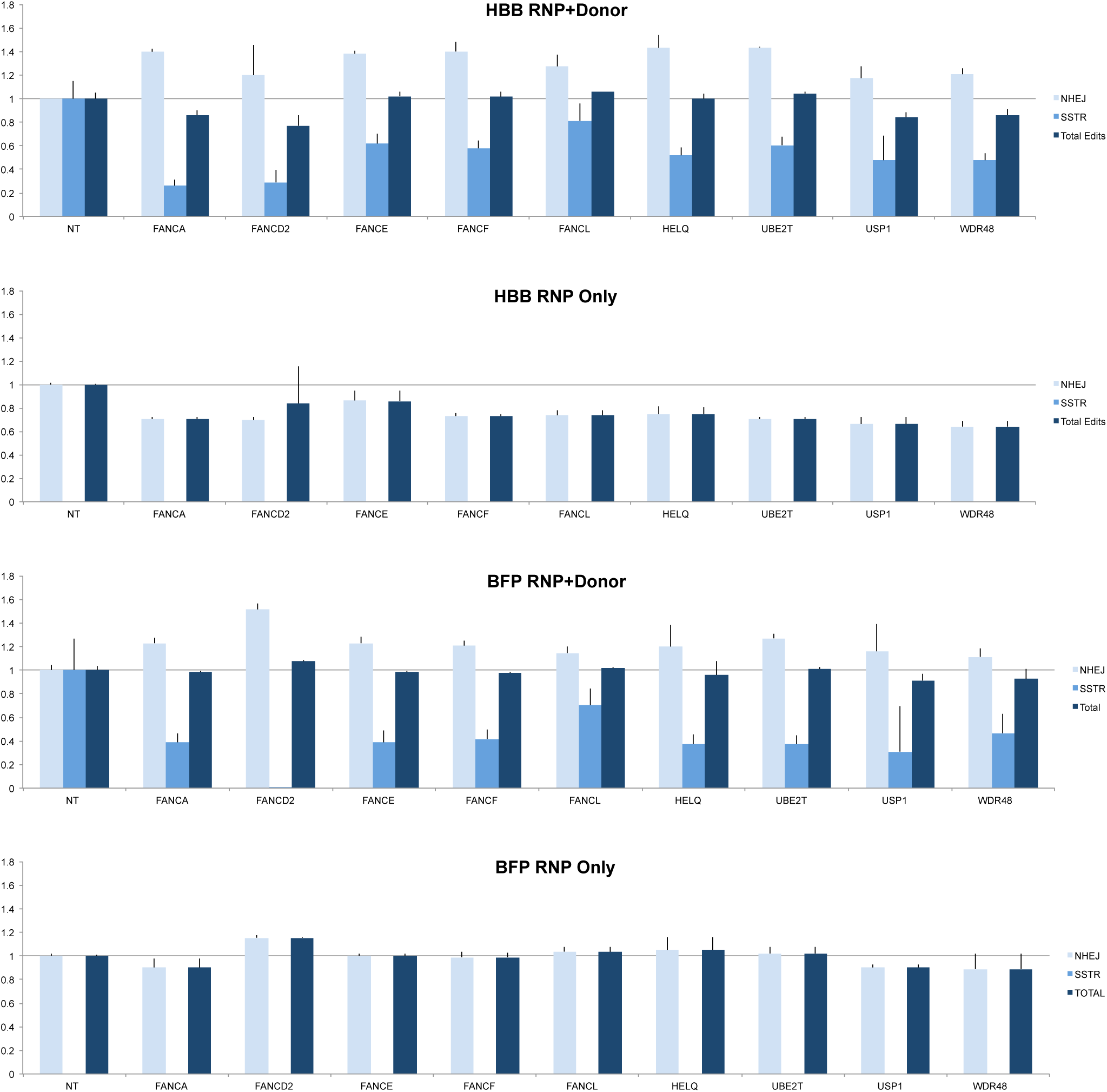
Total editing remains consistent when SSTR is disrupted. Indel (NHEJ, light blue), SSTR (medium blue), and total (dark blue) editing events were quantified for the indicated CRISPRi cell lines and normalized to untargeted controls. **(A)** RNP targeting the HBB locus with donor ssODN. **(B)** RNP targeting the HBB locus without ssODN. **(C)** RNP targeting the BFP locus with donor ssODN. **(D)** RNP targeting the BFP locus without ssODN. Data presented are the mean±sd of biological duplicates.

**Extended Data Figure 10:**
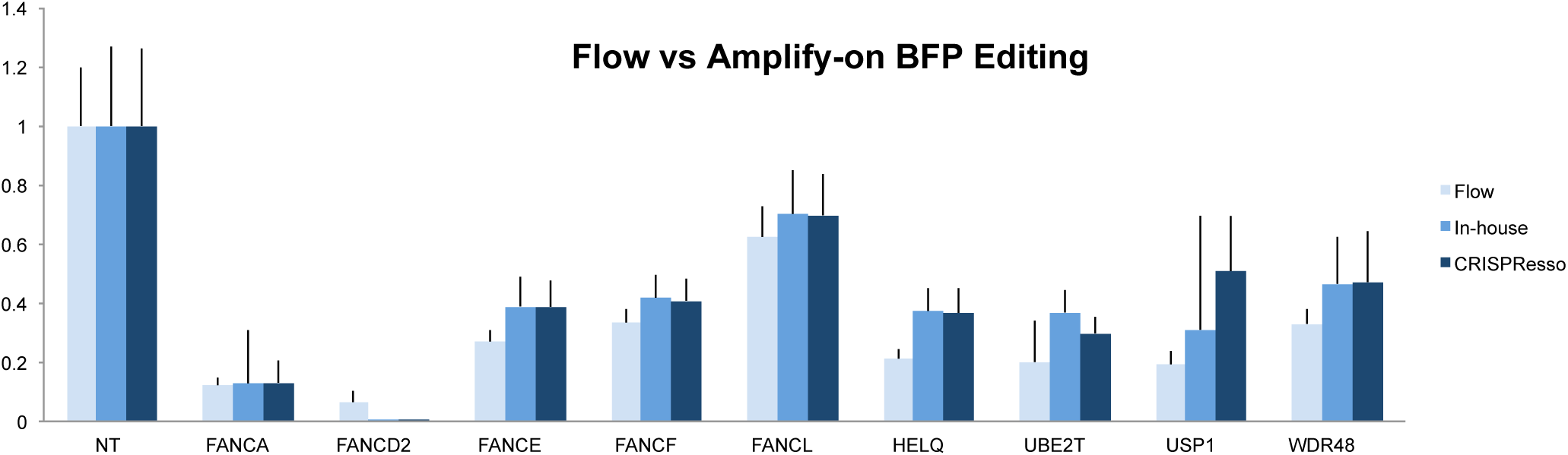
Flow cytometry and amplicon sequencing produce similar editing rates. Editing at the BFP locus was quantified by flow cytometry (light blue), amplicon sequencing followed by in-house bioinformatic analysis (medium blue), or amplicon sequencing followed by CRISPRessoPOOL^33^ analysis (dark blue). Data presented are biological duplicate.

**Document S1:** Materials and Methods (Word doc)

**Document S2:** Molecular biology reagents (Excel doc)

**Document S3:** Processed screen data – collapsed gene scores (Excel doc)

## Acknowledgements

We thank members of the Corn lab for helpful discussions about the manuscript. This work used the Vincent J. Coates Genomics Sequencing Laboratory at UC Berkeley, supported by NIH S10 OD018174 Instrumentation Grant. We thank the Berkeley Macrolab for support with protein expression and purification. This work was supported by grants from the Li Ka Shing Foundation, the Heritage foundation, and the Fanconi Anemia Research Foundation.

## Author Contributions

CDR and JEC designed experiments; CDR, AJS, and SF performed pooled screens; CDR, KRK, and SJF performed follow-up experiments; CDR, NLB, and JEC analyzed data; CDR and JEC wrote the manuscript.

## References

1. DeWitt, M. A. et al. Selection-free genome editing of the sickle mutation in human adult hematopoietic stem/progenitor cells. Sci Transl Med 8, 360ra134–360ra134 (2016).

2. Cox, D. B. T., Platt, R. J. & Zhang, F. Therapeutic genome editing: prospects and challenges. Nat Med 21, 121–131 (2015).

3. Sternberg, S. H. & Doudna, J. A. Expanding the Biologist’s Toolkit with CRISPR-Cas9. Mol Cell 58, 568–574 (2015).

4. Maggio, I. & Gonçalves, M. A. F. V. Genome editing at the crossroads of delivery, specificity, and fidelity. Trends in biotechnology 33, 280–291 (2015).

5. Chen, F. et al. High-frequency genome editing using ssDNA oligonucleotides with zinc-finger nucleases. Nat Meth 8, 753–755 (2011).

6. Jasin, M. Genetic manipulation of genomes with rare-cutting endonucleases. Trends Genet 12, 224– 228 (1996).

7. Richardson, C. D., Ray, G. J., DeWitt, M. A., Curie, G. L. & Corn, J. E. Enhancing homology-directed genome editing by catalytically active and inactive CRISPR-Cas9 using asymmetric donor DNA. Nat Biotechnol (2016). doi:10.1038/nbt.3481

8. Sander, J. D. & Joung, J. K. CRISPR-Cas systems for editing, regulating and targeting genomes. Nat Biotechnol 32, 347–355 (2014).

9. Wright, A. V. et al. Rational design of a split-Cas9 enzyme complex. Proceedings of the National Academy of Sciences 112, 2984–2989 (2015).

10. Kim, S., Kim, D., Cho, S. W., Kim, J. & Kim, J.-S. Highly efficient RNA-guided genome editing in human cells via delivery of purified Cas9 ribonucleoproteins. Genome Research 24, 1012–1019 (2014).

11. Horlbeck, M. A., Gilbert, L. A., Villalta, J. E. & Adamson, B. Compact and highly active next-generation libraries for CRISPR-mediated gene repression and activation. eLife (2016). doi:10.7554/eLife.19760.001

12. Huang, D. W., Sherman, B. T. & Lempicki, R. A. Systematic and integrative analysis of large gene lists using DAVID bioinformatics resources. UNKNOWN 4, 44–57 (2008).

13. George, B. et al. Regulation and function of Myb-binding protein 1A (MYBBP1A) in cellular senescence and pathogenesis of head and neck cancer. Cancer Lett 358, 191–199 (2015).

14. Subramanian, A. et al. Gene set enrichment analysis: a knowledge-based approach for interpreting genome-wide expression profiles. Proc Natl Acad Sci USA 102, 15545–15550 (2005).

15. Lin, S., Staahl, B. T., Alla, R. K. & Doudna, J. A. Enhanced homology-directed human genome engineering by controlled timing of CRISPR/Cas9 delivery. eLife 4, (2014).

16. Bothmer, A. et al. Characterization of the interplay between DNA repair and CRISPR/Cas9-induced DNA lesions at an endogenous locus. Nat Commun 8, 1–12 (2016).

17. Davis, L. & Maizels, N. Homology-directed repair of DNA nicks via pathways distinct from canonical double-strand break repair. Proc Natl Acad Sci USA 111, E924–32 (2014).

18. Yang, Y.-G. et al. The Fanconi anemia group A protein modulates homologous repair of DNA double-strand breaks in mammalian cells. Carcinogenesis 26, 1731–1740 (2005).

19. Hustedt, N. & Durocher, D. The control of DNA repair by the cell cycle. Nat Cell Biol (2017). doi:10.1038/ncb3452

20. Walden, H. & Deans, A. J. The Fanconi anemia DNA repair pathway: structural and functional insights into a complex disorder. Annu Rev Biophys 43, 257–278 (2014).

21. Moldovan, G.-L. & D’Andrea, A. D. How the fanconi anemia pathway guards the genome. Annu Rev Genet 43, 223–249 (2009).

22. Palovcak, A., Liu, W., Yuan, F. & Zhang, Y. Maintenance of genome stability by Fanconi anemia proteins. Cell Biosci 7, 8 (2017).

23. Ellis, N. A., Groden, J., Ye, T. Z., Straughen, J. & Lennon, D. J. The Bloom’s syndrome gene product is homologous to RecQ helicases. Cell 83, 655–666 (1995).

24. Adelman, C. A. et al. HELQ promotes RAD51 paralogue-dependent repair to avert germ cell loss and tumorigenesis. Nature 502, 381–384 (2013).

25. Chun, J., Buechelmaier, E. S. & Powell, S. N. Rad51 paralog complexes BCDX2 and CX3 act at different stages in the BRCA1-BRCA2-dependent homologous recombination pathway. Mol Cell Biol 33, 387–395 (2013).

26. Adamo, A. et al. Preventing nonhomologous end joining suppresses DNA repair defects of Fanconi anemia. Mol Cell 39, 25–35 (2010).

27. Renaud, E., Barascu, A. & Rosselli, F. Impaired TIP60-mediated H4K16 acetylation accounts for the aberrant chromatin accumulation of 53BP1 and RAP80 in Fanconi anemia pathway-deficient cells. Nucleic Acids Research 44, 648–656 (2016).

28. Howard, S. M., Yanez, D. A. & Stark, J. M. DNA damage response factors from diverse pathways, including DNA crosslink repair, mediate alternative end joining. PLoS Genet. 11, e1004943 (2015).

29. van Overbeek, M. et al. DNA Repair Profiling Reveals Nonrandom Outcomes at Cas9-Mediated Breaks. Mol Cell 1–30 (2016). doi:10.1016/j.molcel.2016.06.037

30. Song, J. et al. RS-1 enhances CRISPR/Cas9- and TALEN-mediated knock-in efficiency. Nat Commun 7, 10548 (2016).

31. Orthwein, A. et al. A mechanism for the suppression of homologous recombination in G1 cells. Nature (2015). doi:10.1038/nature16142

32. Knight, S. C. et al. Dynamics of CRISPR-Cas9 genome interrogation in living cells. 350, 823–826 (2015).

33. Pinello, L. et al. Analyzing CRISPR genome-editing experiments with CRISPResso. Nat Biotechnol 34, 695–697 (2016).

